# Yeast synthetic minimal biosensors for evaluating protein production

**DOI:** 10.1101/2020.11.03.367615

**Authors:** Kai Peng, Heinrich Koukamp, Isak S. Pretorius, Ian T. Paulsen

**Affiliations:** ARC Centre of Excellence in Synthetic Biology, Department of Molecular Sciences, Macquarie University, NSW 2109, Australia; Biomolecular Discovery and Design Research Centre, Macquarie University, Sydney, NSW 2109, Australia; Chancellery, Macquarie University, Sydney, NSW 2109, Australia

**Keywords:** yeast, unfolded protein response, biosensor, minimal promoter, secretion

## Abstract

The unfolded protein response (UPR) is a highly conserved cellular response in eukaryotic cells to counteract endoplasmic reticulum (ER) stress, typically triggered by unfolded protein accumulation. In addition to its relevance to human diseases like cancer cell development, the induction of the UPR has a significant impact on recombinant protein production yields in microbial cell factories, including the industrial workhorse *Saccharomyces cerevisiae.* Being able to accurately detect and measure this ER stress response in single cells, enables the rapid optimisation of protein production conditions and high-throughput strain selection strategies. Current methodologies to monitor the UPR in *S. cerevisiae* are often temporally and spatially removed from the cultivation stage, or lack updated systematic evaluation. To this end we constructed and systematically evaluated a series of high-throughput UPR sensors by different designs, incorporating either yeast native UPR promoters or novel synthetic minimal UPR promoters. The native promoters of *DER1* and *ERO1* were identified to have suitable UPR biosensor properties and served as an expression level guide for orthogonal sensor benchmarking. Our best synthetic minimal sensor, SM1, was only 98 bp in length, had minimal homology to other native yeast sequences and displayed superior sensor characteristics. Using this synthetic minimal UPR sensor, we demonstrate its ability to accurately discriminate between cells expressing different heterologous proteins and at varying production levels. Our sensor is thus a novel high-throughput tool for determining expression/engineering strategies for optimal heterologous protein production.

## Introduction

The unfolded protein response (UPR) is a highly conserved cellular response in eukaryotic cells to maintain the ER folding homeostasis and acts as a protective buffer against excessive endoplasmic reticulum (ER) stress (Mori 2009, Walter 2011). When the protein folding requirements in the ER exceeds the capacity of the ER folding machinery, the accumulation of misfolded proteins typically results in the UPR activation. Disruptions in the ER membrane morphology or lipid composition have also been shown to trigger the UPR (Lajoie 2012, Promlek 2013, Volmer 2013, Volmer 2015). The UPR relieves ER stress and restores homeostasis by modulating the transcription of UPR responsive genes (UPR genes) (Kimata 2006) involved in processes to modulate protein translocation rates in-and-out of the ER, fine-tune protein chaperone level, expand the volume of the ER (Bernale 2006) and enhance the ER associated degradation (ERAD) of terminally misfolded proteins (Travers 2000, Ng 2000, Hou 2012, Kimata 2006). In higher eukaryotes, the UPR also slows the cellular protein synthesis rates through translational control and mRNA decay, while constitutively activated UPR has been proposed to lead to cell apoptosis (Walter 2011). The absence or malfunction of the UPR has been linked to various human diseases, including retinitis pigmentosa, multiple myeloma, severe obesity, enveloped virus infections and many types of cancer (Ma 2004, Walter 2011, Volmer 2015). The importance of this ER quality control mechanism is highlighted in mammalian cells by the presence of multiple UPR pathways, namely the PERK, ATF6 and IRE1 pathways, allowing precise control of metabolism, translation and apoptosis to resolve the ER stress (Lin 2007, Walter 2011).

In the unicellular eukaryote *Saccharomyces cerevisiae*, the UPR pathway is mono-threaded, involving two major components: (1) the ER membrane protein Ire1p that acts as ER stress sensor and the signal transducer, (2) and the leucine-zipper transcription activator Hac1p (homolog of the mammalian Xbp1p) (Cox 1996) as the UPR effector. Hac1p regulates the expression of approximately 400 UPR genes, accounting for approximately 6% of genes in the yeast genome (Travers 2000, Kimata 2006). At the onset of ER stress, the Ire1p senses and binds the misfolded proteins via its ER luminal misfolded protein binding domain (Pincus 2010, Gardner 2011), which then triggers the higher-order oligomerization of Ire1p (Korennykh 2009, van Anken 2014). This process is attenuated through the interaction between Ire1p, the essential chaperone Kar2p and unfolded protein in the ER lumen (Pincus 2010). The clustering of Ire1p subsequently activates the RNase activity of its cytosolic domains allowing the spliceosome-independent removal of the 252bp intron from immature *HAC1* mRNA (Cox & Walter 1996; Sidrauski 1997). The mature *HAC1* mRNA produced via this alternative splicing is translated into Hac1p, which subsequently gets imported into the nucleus where it binds to upstream activating sequence, called UPR element (UPRE), to regulate UPR gene expression (Mori 1992, Cox 1996, Mori 1998, Fordyce 2012).

The relatively simple UPR pathway in *S. cerevisiae* has served as a model for elucidating the evolution of the UPR in eukaryotes, the mechanisms of unfolded protein detection, signal transduction mechanism and the transcriptional response, which ultimately restores folding homeostasis. *S. cerevisiae* is also an important cell factory for both recombinant protein and metabolite production (Hou 2012, Gasser 2008, Xu 2014). And engineering strategies often involve the overexpression of native and heterologous proteins, frequently overwhelming the ER’s folding capacity and activating the UPR, which has a direct and usually negative impact on protein production titre. Therefore, in addition to aiding our fundamental understanding of the UPR, the real-time detection and measurement of the UPR in *S. cerevisiae* offers valuable insights into a cell factory’s capability to correctly process protein products and allows for high-throughput optimisation of cultivation and protein expression strategies.

Conventional methods for detecting UPR induction relies on the detection of the spliced form of *HAC1* mRNA (Cox 1996, Merksamer 2008), and an estimate of the activation intensity determined from the ratio of spliced to unspliced levels of *HAC1* mRNA (Pincus 2010, Le 2016). These RNA-based methodologies are time consuming and susceptible to large technical errors due to the many steps involved. The destructive nature of these methods does not allow for high-throughput isolation of individual cells of interest. More recently, biological sensors have been developed for mechanistic studies into UPR regulation. Early sensors exploited the Hac1p dependent UPRE of UPR genes, with constructs consisting of one or more copies of 22-bp redundant UPRE consensus sequence 1 (rUPRE-1, Mori 1992; later refined to the CAGCGTG core sequence, Mori 1998), placed upstream of a *CYC1* core promoter and used enzymatic reporters like *β*-galactosidase (Mori 1992, Cox 1996, Travers 2000, Patil 2004), which are now mostly superseded by fluorescent proteins (Travers 2000, Xu 2005, Merksamer 2008). In addition to the UPR transcription mechanism, several UPR sensor designs also tapped into other UPR related non-transcriptional cellular process as indirect reflection of the UPR, like using redox-sensitive GFP (eroGFP) to monitor the redox change in the ER lumen caused by UPR activation (Merksamer 2008, Lajoie 2012); and making UPR-specific alternative splicing reporter by substituting GFP for the first exon of the *HAC1* gene, that only produces GFP when the alternative splicing occurs (Aragon et al, 2009, Pincus 2010, Lajoie 2012).

The previous UPR transcription-based sensors have relied upon either the 22bp redundant UPRE1 (Mori 1992) plus native non-UPR promoter like *CYC1*; or upon limited choices of native UPR promoters like that of *HAC1*, *KAR2* (Cedras 2020) and *ERO1* (Patil 2004, Pincus 2014), that could be subjected to additional non-UPR specific regulation. While in the last decade substantial advances have been made in both UPRE (Fordyce et al 2012) and synthetic minimal promoter in *S. cerevisiae* (Redden et al 2015). In the work by Fordyce et al. (2012), the authors used systematically designed DNA fragments to investigated their affinity to the Hac1p binding *in vitro*, and were able to subsequently refine the previously reported Hac1p binding site to a concise 12 bp core UPRE1 (represented by GGACAGCGTGTC, UPRE1 hereafter) and 10bp core UPRE2 (represented by CTACGTGTCT, UPRE2 hereafter). This resolved some of the ambiguity remained about the prevalence of UPREs in UPR responsive genes, where it was previously thought that fewer than half of all the UPR genes contained UPRE (Travers 2000, Patil 2004). The harmonization of UPRE also absorbed the previously termed UPRE3 (Patil 2004) into the new framework. On the other hand, refinement of functional core promoters has been reported in *S. cerevisiae* (Redden et al. 2015), demonstrating low but constitutive expression of genes from promoters of merely about 100bp. These minimal promoters can be easily converted into strong inducible promoters through the incorporation of upstream activation sequences (UAS).

These recent developments have prompted the updated synthetic minimal UPR sensor design described in this study. Several minimal sensors were designed, characterised and systematically evaluated against sensors employing native UPR promoters. We selected the G-SM1 sensor as the construct displaying superior signal resolution, dynamic range and signal strength. Furthermore, we show the this synthetic minimal UPR biosensor could be used to distinguish between cells with different heterologous protein secretion levels, emphasising the potential of these biosensors as qualitative high-throughput screening tools for determining optimal expression/engineering strategies for producing secretory heterologous proteins.

## Results and Discussions

### Characterisation of native UPR gene expression profiles

We first investigated if and which native UPR promoters could be effective synthetic parts in building UPR sensors, which also helps to establish a more holistic view of the expression characteristics of the native UPR genes in baker’s yeast. Except for some commonly used UPR genes like *KAR2*, *ERO1* and *HAC1* (Cedras 2020, Pincus 2014), the transcriptional activation profiles of many native UPR promoters have not been documented. To this end we selected and characterised the expression profiles of 9 putative promoters of known UPR genes in *S. cerevisiae* (Travers 2000), covering different functional categories of the secretory pathway: *DER1* for misfolded protein degradation, *ERO1*, *EUG1*, *KAR2* and *PDI1* for polypeptide folding, *PMT1* for protein modification, *SEC12* and *SEC62* for protein trafficking, plus *HAC1* the UPR effector. As UPR genes, their promoters were expected to contain UPRE sequences, upstream of a functional core promoter. In addition to the known UPRE of *KAR2* and *ERO1*, the predicted UPREs the other genes were listed in Table 1.

**Table 1.**
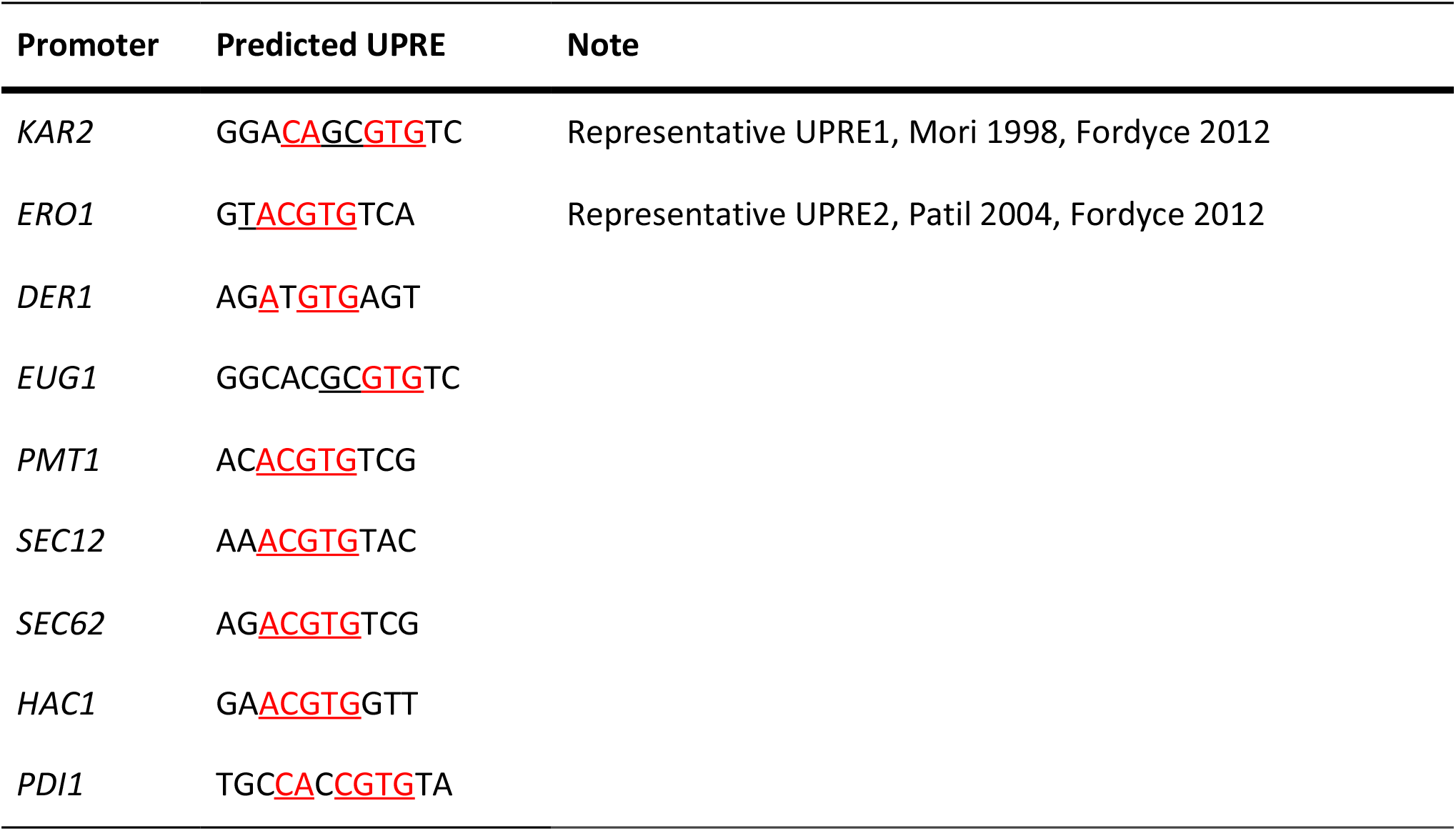
The UPRE native UPR promoters evaluated in this study.

The UPR biosensor-containing cells were challenged by increasing concentrations of tunicamycin (Tm), which represented increasing ER stress levels. Tunicamycin is a commonly used UPR chemical inducer and promotes the aberrant folding of proteins by inhibiting *N*-linked glycosylation of newly synthesized polypeptides within the ER lumen (Travers 2000). Using the eGFP fluorescence output of each native sensor, we evaluated its ability to resolve differences in Tm concentrations (resolution), the ratio between its maximal and basal response signals (dynamic range) and its absolute fluorescence level (AFU, signal intensity).

All the nine constructs had higher unchallenged fluorescence levels than the P-Gal1 control, indicating basal level expression in the absence of ER stress (Figure 1). This basal expression of UPR genes was expected, since these gene products are part of the native (unstressed) protein processing and recycling pathways. The P-ERO1, P-HAC1, P-KAR2 and P-PDI1 sensors had the highest basal expression levels (the high AFU group, Figure 1b), with P-KAR2 being the strongest showing over 65-fold higher AFU compared to that of P-Gal1; while the expression from P-DER1, P-EUG1, P-PMT1, P-SEC12 and P-SEC62 were significantly lower (low AFU group, Figure 1a), ranging from 2- to 10-folds of the P-Gal1 level. When induced, the signal of all nine constructs increased with the rise in the Tm concentration. The P-DER1, P-ERO1, P-HAC1 and P-SEC12 sensors had the best sensitivity to low stress, responding to Tm concentration of 0.2 μg/mL, while the signal of all the constructs plateaued after the concentration of 1 μg/mL of Tm (Figure 1). This signal ceiling could signify the maximal response at which the Ire1p-Hac1p mediated UPR signalling pathway has reached saturation in its transcription activation activity or alternatively, it could represent an unfolded protein stress level at which protein synthesis including the eGFP reporter itself is inhibited.

**Figure 1.**
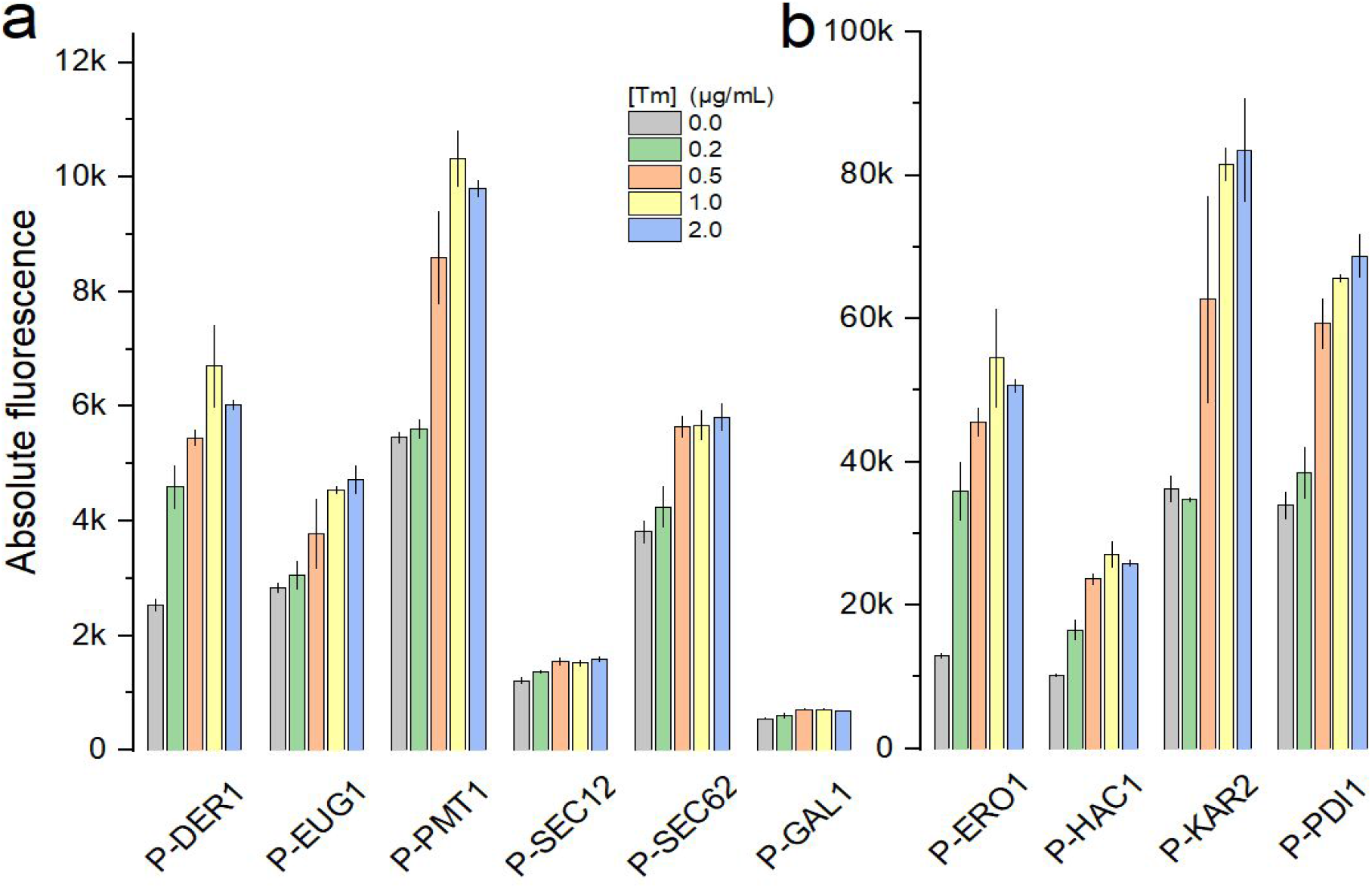
The response of UPR sensors constructed with native UPR promoters that had relatively low (a) and high (b) absolute fluorescence intensity (AFU). Cells were incubated in tunicamycin concentration gradient for 4 hours. Each point represented 3 biological replicates. The *GAL1* promoter was used as non-UPR control.

**Figure 2.**
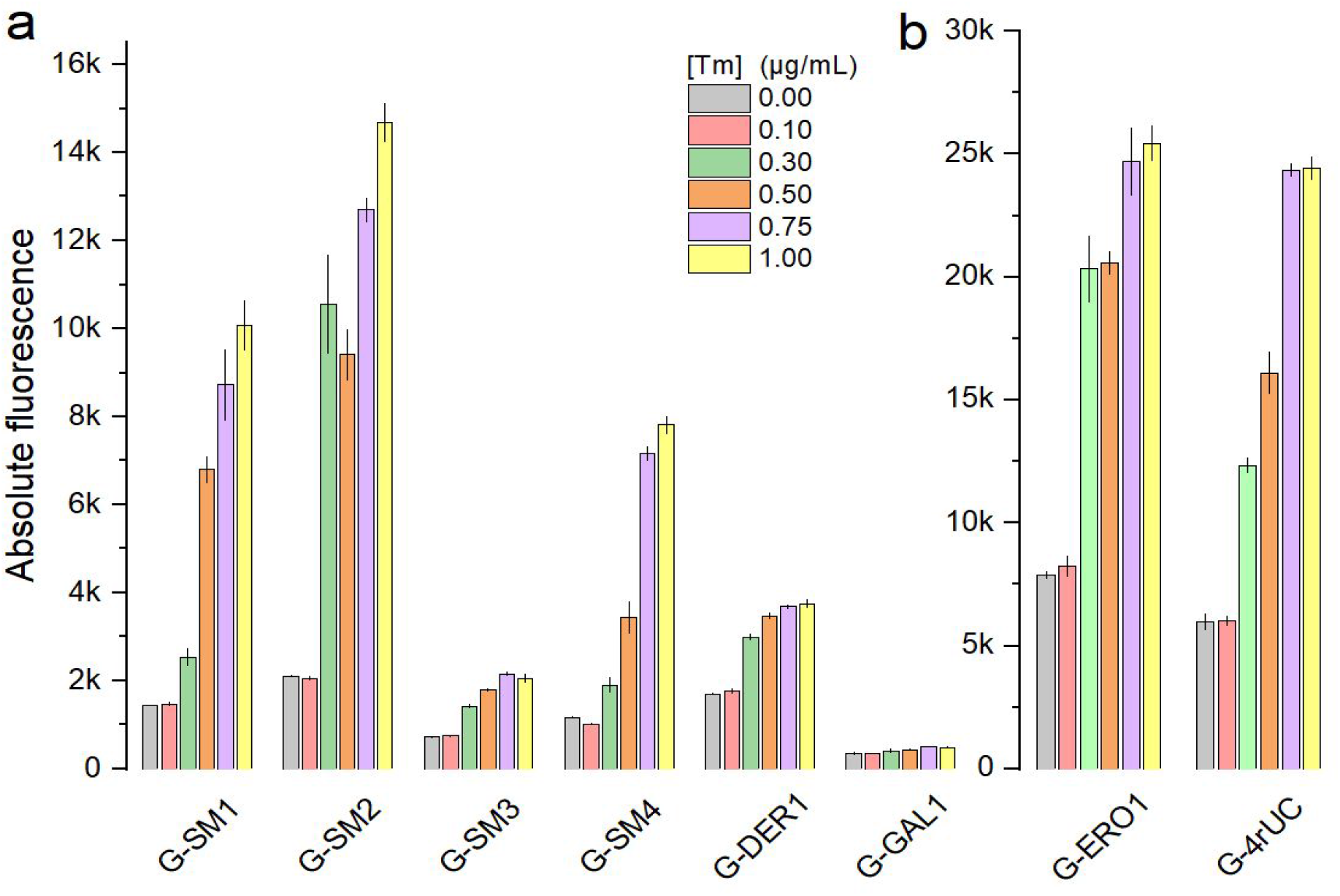
The response of UPR sensor constructed with synthetic minimal, semi-synthetic, DER1 and ERO1 UPR promoters. The sensors were divided into the relatively low (a) and high (b) AFU groups for display. Cells were incubated in tunicamycin concentration gradient for 6 hours. Each point represented at least 3 biological replicates. The *GAL1* promoter was used as non-UPR control.

For constructs with fluorescence in the lower range, the P-DER1 had the best working range and the biggest dynamic range, resolving 0, 0.2, 0.5 and 1.0 μg/mL reaching a maximal signal of 2.6 times the basal fluorescence level. In the high AFU group, the P-ERO1 had the best dynamic range of 4.2 times of its basal fluorescence, and along with the P-HAC1 sensor achieved the same resolution as P-DER1. Although P-HAC1 shared these favourable characteristics when evaluated with Tm stress, it was unable to separate stress induced by another UPR inducing agent, dithiothreitol (DTT) at 2mM and 3mM (Figure S1). Displaying superior dynamic and working ranges compared to the promoters in the low/high AFU group, we selected P-DER1 and P-ERO1 as the most suitable representative native-promoter UPR sensors for benchmarking purposes.

### Construction and Evaluation of Synthetic Minimal UPR Sensors

In the current era of synthetic biology, researchers seek to extract genetic components into modular building blocks to make synthetic functional parts or circuits. This ideology, together with the latest developments in *S. cerevisiae* UPRE refinement (Fordyce 2012) and synthetic minimal promoter methodology (Redden 2015) inspired the design of the synthetic minimal UPR sensors described here. In addition to a more concise design embracing “nucleotide economics”, a key consideration was to limit the amount of native DNA sequences to improve long term stability of the construct and reduce unwanted activation by cryptic transcription activation sequence. To this end, four synthetic minimal UPR promoters were constructed, containing one or four copies of concise 12bp UPRE1 (GGACAGCGTGTC) or 10bp UPRE2 (CTACGTGTCA) upstream of a synthetic minimal core promoter (Redden 2015). Similar to the native biosensor cassettes, these synthetic minimal promoters were used to drive the expression of eGFP as reporter. Prior studies have made use of a native constitutive promoter *CYC1* with four copies of the 22bp redundant UPRE1 placed upstream (henceforth referred to as the ‘semi-synthetic’ sensor construct) (Merksamer 2008, Pincus 2010, De Ruijter 2018). We then compared the UPR stress response characteristics of each category, namely the native UPR promoters, the synthetic minimal promoters and the semi-synthetic promoter, in order to determine the ability of each to report on UPR stress. All of these sensors were chromosomally integrated as a single copy at the *flo8* locus. This was done to achieve enhanced homogeneity (reduced signal noise) of single cell fluorescence within the population (Figure S2), due to reduced construct copy number variability and enhanced construct stability. We repeated the Tm gradient stress test for all of these constructs and evaluated the sensor response based on the criteria of resolution, dynamic range and signal intensity as the previous section. Since high level of heterologous protein production is a known activator of the UPR, consequently it is possible that eGFP protein synthesis itself may exert stress or compete for ER folding machinery, the maximal GFP signal was also considered as evaluation criterium.

The G-DER1 and G-SM3 sensors were the only ones able to significantly resolve UPR induction difference between unstressed and Tm concentration of 0.1 μg/mL (p<0.05). Since both *DER1* and *ERO1* was selected based on their ability to display significant activation at a 0.2μg/mL Tm, this lower stress level may signify the turning point where significant unfolded protein stress had accumulated to initiate the UPR. Except for G-ERO1 and G-SM2, all sensors could distinguish between 0.1, 0.3, 0.5 and 0.75 μg/mL Tm (p<0.05).

At the higher stress levels only the synthetic sensors, G-M1G, G-M2G and G-M4G successfully resolved 0.75μg/mL from 1μg/mL (p<0.05). The G-M1G and G-M4G thus had the widest working range of all the sensor constructs, performing well between 0.1-1 μg/mL Tm. Although the chimeric sensor had a distinct separation between Tm concentrations of 0.1 - 0.75 μg/mL, it lacked sensitivity both at the lower and higher stress levels.

For single cell application, sensors are subjected to the inherent stochastic biological variability between cells, and sensors with a large signal dynamic range are highly desired. Here again, the G-SM1, G-SM2 and G-SM4 sensors had the largest dynamic ranges of ~7 fold that of their respective uninduced fluorescence levels. Although G-DER1 was sensitive to 0.1 μg/mL Tm, it had the lowest dynamic range value of all the evaluated sensors, only managing a ~2 fold increase.

Overall, the G-SM1 and G-SM4 synthetic minimal UPR sensors had the best dynamic signal and working ranges. In addition, they had intermediate signal strength throughout their working ranges, reducing concerns about the influence of eGFP production. The UPRE2 elicited weaker transcriptional activation and even with four copies, the G-SM4 only had ~80% the signal strength of G-SM1 with single UPRE1.

### Evaluation of HAC1-independent biosensor noise

Besides the resolution, dynamic range and signal intensity criteria, we were also interested in determining whether the biosensor exclusively responds to Hac1p-transcriptional activation in our cultivation conditions. Complete orthogonality is not always achievable in biological systems, considering the complex and interconnected nature of cellular metabolism; however, with careful biosensor design we aimed to minimise any indirect off-target responses or noise. While our bioinformatic analysis of the synthetic minimal promoter constructs did not reveal any known stress responsive UAS elements other than the UPREs, we wanted to assess whether the sensor basal fluorescence levels were influenced by transient Hac1p transcription factor levels under unstressed conditions. Since Hac1p is known to be the sole activator of UPR in yeast, the Hac1p-dependent and Hac1p-independent signal to noise ratios were determined in UPR stressed wild type and *hac1* deleted strains (Figure 3). The P-DER1, P-ERO1, P-4rUC, P-SM1 and P-SM4 constructs were chosen to represent the native, semi-synthetic and synthetic minimal UPR promoter designs.

**Figure 3.**
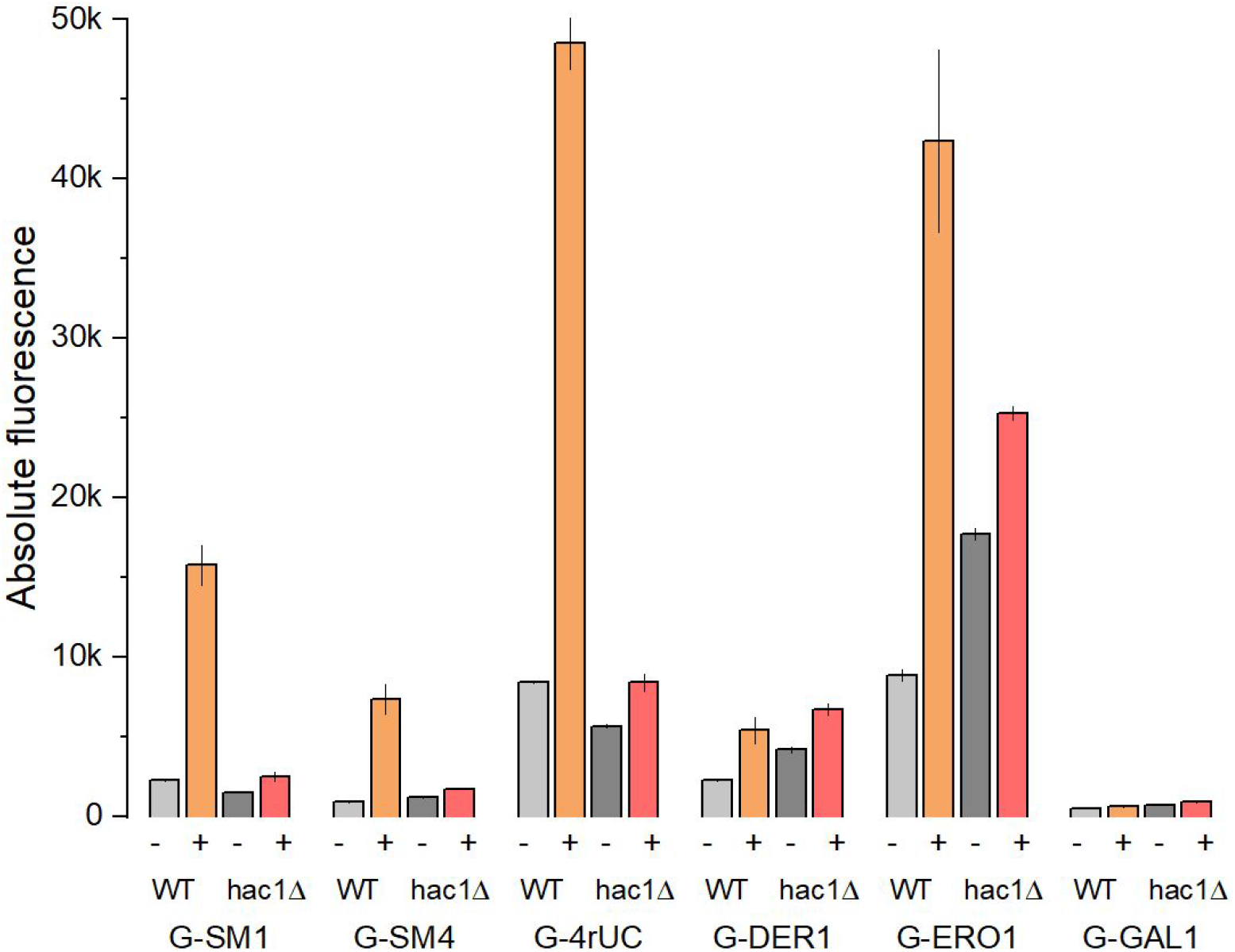
The fluorescence output of UPR sensors in wildtype and *hac1* deleted strains. The unfolded protein response was activated through treatment with the presence (+) or absence (−) of 0.5 μg/mL tunicamycin for 6 hours. The *GAL1* promoter served as a control. The error bars represent the standard deviation between biological quadruplicates.

For all the five UPR sensors, unstressed basal-level fluorescence was observed in both *hac1Δ* and wildtype strains over the P-GAL1 control. The P-4rUC sensor had the highest activation due to basal (unstressed) levels of Hac1p, losing 26% of its signal strength when *HAC1* was deleted. Unlike the native P-ERO1 and P-DER1 promoter sensor, the P-SM1, P-SM2 and P-4rUC sensors in Tm challenged *hac1Δ* deleted strains had similar signal strengths compared to their corresponding wild type counterparts under unchallenged conditions. This would suggest that under our cultivation conditions, the synthetic minimal and chimeric sensor P-4rUC are exclusively stimulated by the abundance of Hac1p transcription factor. The activation of the native sensors in the *hac1Δ* strain, when not subjected to Tm stress was unexpected, but demonstrates inherent safety circuitry (or evolutionary redundancy) within biological systems to protect itself.

The relative unresponsiveness of the synthetic minimal promoters to ER stress when *HAC1* is deleted demonstrates their dependence on the functional Hac1p transcriptional activation, implying isolation from other non-specific or indirect activation by unfolded protein accumulation. Although both the SM1 and SM4 synthetic minimal promoters had superior dynamic and working ranges, we selected the SM1 for the subsequent protein production stress work, as it was the smallest of the synthetic sensors at only 98 bp compared to 126 bp of SM4.

#### Detecting Folding Stress Caused by Misfolded Proteins

*S. cerevisiae* serves as a model organism to study the mechanisms of the UPR, while also being a persistent protein cell factory organism of choice. This has been frequently noted that, in addition to aberrant protein folding or aggregation, the overproduction of protein itself activates the UPR. This implies that all the standard methods of increasing protein production, such as increasing gene of interest copy number and the use of strong promoters to drive gene expression could all ultimately stimulate the UPR. Although the initial increased ER folding capacity brought by the UPR may benefit some protein products, continuous high-level UPR activation generally often results in a negative net impact on the secretion titres (Ilmén 2011). For this reason, a quantitative evaluation and early detection system of ER stress in protein producing cell factories is of great importance; and in addition, could serve as a high-throughput tool to understand strain-dependent or protein-specific factors contributing to UPR induction.

To this end, we first evaluated our synthetic minimal sensor’s ability to resolve ER stress caused by protein production stresses, we overexpressed carboxypeptidase Y (CPY) and its mutated version (CPY*) in our SM1 biosensor strain (Figure 4). CPY is a *S. cerevisiae* native vacuolar peptidase, while the mutant CPY* version has a single amino acid mutation of G255R, which makes it prone to aberrant folding and aggregation in the ER lumen. CPY* expression is frequently used as a model system to stimulate UPR activation in yeast (Finger1993, Ng 2000, Merksamer 2008).

**Figure 4.**
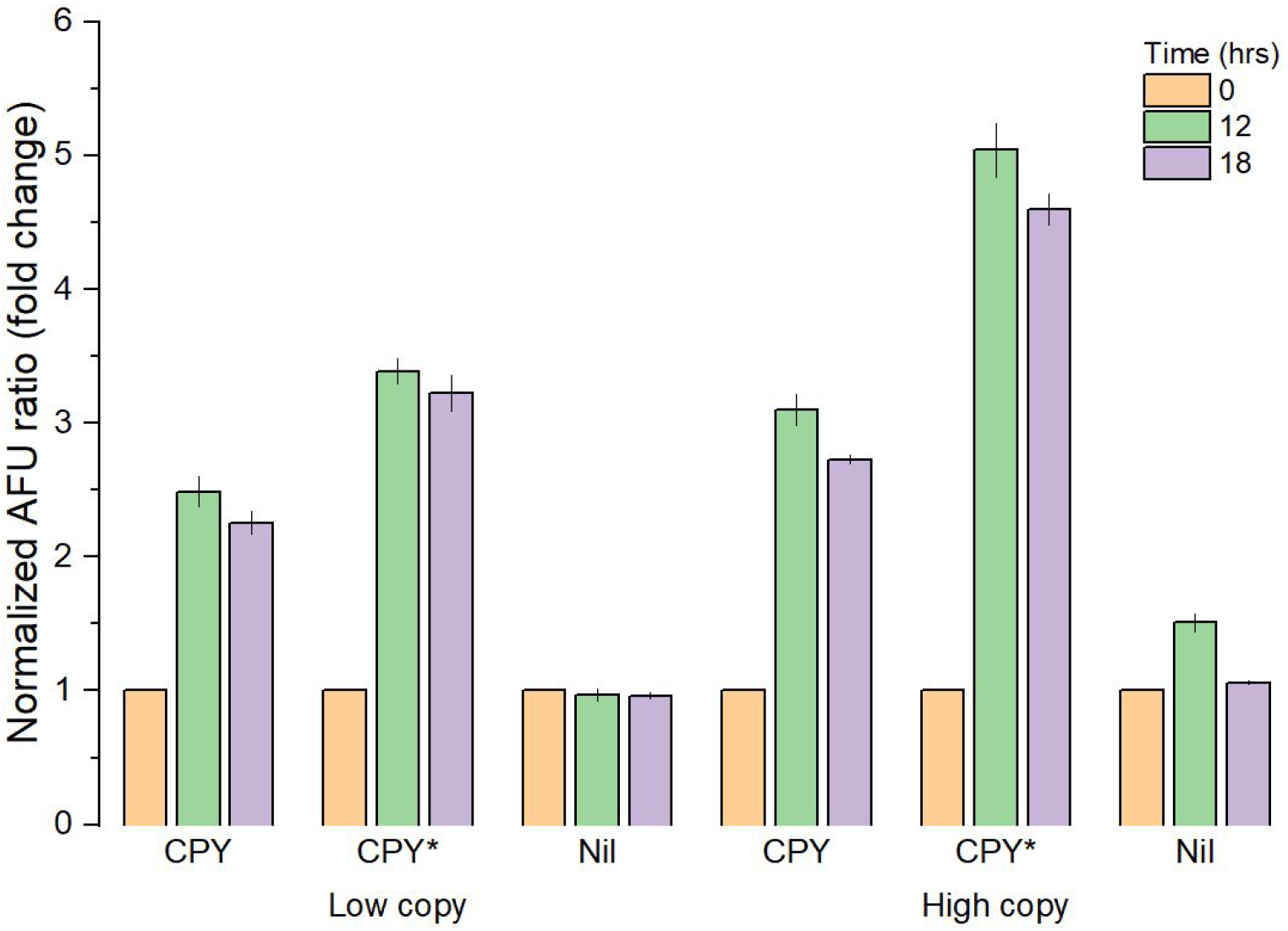
The G-SM1 UPR sensor response in BY4742 producing native CPY and its constantly misfolded form CPY*, via low and high copy number expression vectors. The expression was induced using 2% galactose, and the AFU was measured at 0, 12 and 18 hours. Strain with empty vector were used as control. The AFU was normalized against the time point zero value of each sample, to represent the fold change in signal. The error bars represent the standard deviation from at least three biological replicates.

After 12 hours’ incubation, the overexpression of wild type CPY from both low and high copy (yeast centromeric and 2 micron) vectors, elicited an unfolded protein response of 2.5-fold (P<0.05) and 3-fold (P<0.05) over the basal level at time point zero, respectively (Figure 4). This suggested, in addition to producing mutant forms of native proteins that native proteins, when overexpressed, could cause significant ER stress. This should be taken into account when modulating native host pathways through promoter exchange. Interestingly, the copy number of the CPY expression vector did not make a significant difference to the UPR response (p_low/high_<0.05). It could be speculated that this might indicate a buffering effect at which this native protein can be effectively processed within a set protein load window. On the other hand, the mutated CPY* strains had significantly stronger fluorescence signals than their corresponding CPY strain, with ~3.4- and ~5.1-times of the basal level signal at 12 hours (Figure 4). This finding aligned with previous studies, and thus verified the G-SM1 sensor’s ability to detect this misfolding stress. It was also able to distinguish between strains expressing CPY* from low and high copy vectors, with a ~0.5-fold increase in signal, suggesting that UPR magnitude, for CPY* at least, was a consequence of both the folding difficulty and protein workload. These data demonstrated the G-SM1 sensor could readily detect and measure UPR caused by protein expression levels and known folding difficulty, even for proteins destined for an intracellular compartment.

#### Detecting Folding Stress Caused by Different Heterologous Proteins

One of the key benefits of using yeast as a cell factory, is its ability to efficiently secrete proteins into its environment. We have demonstrated the capacity of our G-SM1 sensor to differentiate between a native and aberrantly folded proteins which only partially completes the journey to the cell membrane. In a study by Ilmén et al. 2011, it was observed that *Trichoderma reesei* Cel7A (Tr.CBH) and the *Talaromyces emersonii* Cel7A (Te.CBH) expressed in yeast, had the lowest and highest secreted active enzyme yields, respectively, out of 14 evaluated cellobiohydrolase I (CBH) enzymes. To evaluate the sensor’s capacity to detect potential UPR induction in strains producing commercially relevant proteins, and whether differences can be detected between different recombinant proteins and production loads, we expressed these two cellobiohydrolase I proteins using low and high copy expression vectors.

Six hours after inducing CBH expression, significant increases in fluorescence of 1.5-fold and 1.8-fold that at T_0_, were observed for strains expressing Tr.CBH on low and high copy vectors respectively (Figure 5a). This protein production stress was detected in these strains, prior to any Tr.CBH activity could be measured in the supernatant (data not shown). This early stimulation of the UPR was not detected in the Te.CBH expressing strains. But at 12 and 18 hours after CBH induction, all strains had increased fluorescence values. At 12 hours after induction, the low and high copy Tr.CBH strains had more than triple the fluorescence, reaching ~3.1 and ~4.6-times their uninduced levels respectively. Similar to the CPY* expression experiment, the high copy Tr.CBH induced significantly stronger signal than the low copy strain, showing increasing protein workload further aggravated the ER stress. By 18 hours, the signal intensity dropped to around ~2.5 and ~4.1-times that of the basal level, which might reflect the cellular effort to re-establish the equilibrium between protein folding capacity and the ER stress. Both the low and high copy Te.CBH expressing strains showed relatively low UPR induction relative to the control strains, with the high-copy strain having a 0.7-fold and the low-copy strain a 0.4-fold increase in fluorescence over their uninduced states (Figure 5a). Our detected biosensor results were consistent with the findings by Ilmén et al (2011), which observed significantly higher levels of spliced *HAC1* mRNA (thus UPR activation) in strains expressing the Tr.CBH than Te.CBH.

**Figure 5.**
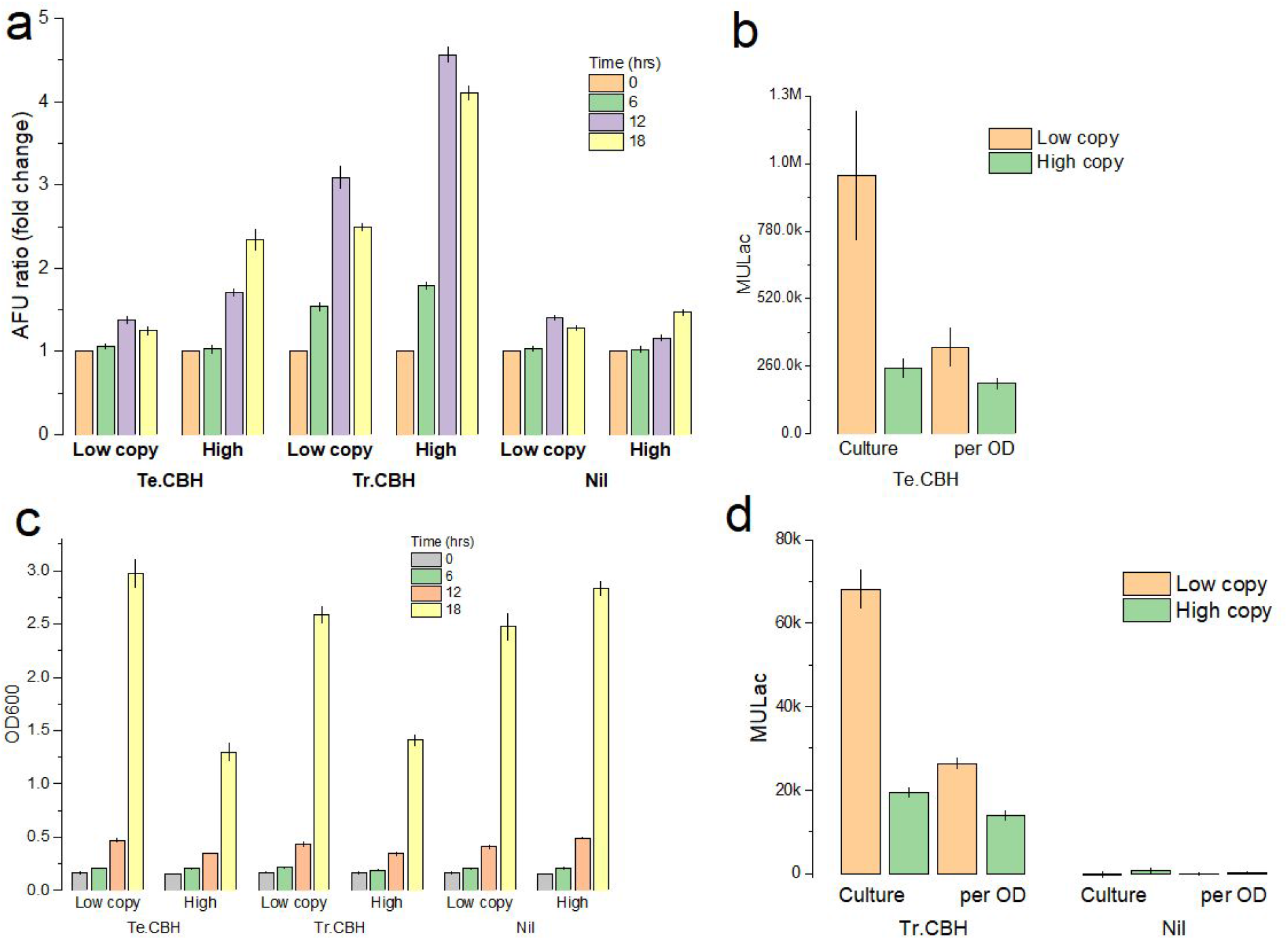
**(a)** The G-SM1 UPR sensor response in BY4742 strains producing Te.CBH and Tr.CBH, via low and high copy expression vectors. The expression CBH was induced using 2% galactose, and the AFU was measured at 0, 6, 12 and 18 hours. Strain with empty vectors were used as control. The AFU was normalized against the time point zero value of each sample, to represent the fold change in signal. (b). The cell density (OD600) of BY4742-M1G strains producing Te.CBH and Tr.CBH, at different time points after galactose induction. **(c, d)** The enzymatic activity of Te.CBH, Tr.CBH and empty vector control measured through MULac assay, from the supernatant of relevant strains after 18 hours of galactose induction. To compare the CBH secretion at per cell level, the MULac readings were also normalized against the cell density (OD600). The error bars represent the sample standard deviation from at least three biological replicates.

For both the Te.CBH and Tr.CBH, the strains harbouring the low copy vectors had significantly higher secreted protein yields (Figure 5c, 5d), as represented through supernatant enzyme activity, than their high-copy counterparts after 18 hours. This correlation between a greater UPR and a concomitant lowered protein secretion illustrates the ability of our biosensor to detect UPR stress before the protein of interest is detected, or the potential use with secreted proteins without simple quantification assays. The smaller difference between the UPR signal of the low and high copy Te.CBH producing strains, was reflected by a smaller difference in secreted protein yields between the same strains, and the larger fluorescence differences seen for the Tr.CBH strains was due to larger difference in secreted product levels. This observation was true even when culture cell densities were taken into account (Figure 5c, 5d).

The two different high-copy vector cellobiohydrolase expressing strains reached similar optical density values after 18 hours (p>0.95). But unlike the low-copy versions, they had significantly impacted cell growth, only reaching around 55% of the low-copy control strain optical density (Figure 5b). Lowered biomass yield is not uncommon in yeast strains overproducing heterologous proteins (Gorgens 2001), and has been reported for Te.CBH expressed from a high-copy vector (Kroukamp et al 2017). Similar reductions in biomass have previously been reported as a direct result of constitutive UPR induction (Valkonen et al 2003), while others suggested that this is a consequence of the increased metabolic burden exerted by high levels of protein synthesis. Our results support the latter observation, since the culture densities in both high-copy strains were low, regardless of the UPR signal strength differences seen between the Te.CBH and Tr.CBH expressing strains. This also suggests that changes in cell biomass are not a suitable proxy to infer general protein production levels or an indication of the UPR induction level.

#### Effect of the strain genetic background on sensor output

We have thus far shown that our biosensor is capable of distinguishing between high and low protein production levels conferred by different vector systems. In addition to expression strategy, protein production is highly dependent on the strain background used and big differences in secretion capacity exist even between laboratory strains (Kroukamp et al 2017). Although the expression levels of many native factors have been shown to have a direct impact on a strain’s protein production and secretion capacity (Kroukamp et al 2018), the actual achievable protein yields are highly protein dependent (Kroukamp et al 2013). Considering the direct impact the UPR has on protein production, and how the strain’s genetic background can modulate this response, we set out to determine whether an elevated UPR would cause similar diminished secreted protein yields in *S. cerevisiae* from a different strain lineage.

Therefore, we investigated how the commonly used *S. cerevisiae* CEN.PK2-1C strain performed when challenged with protein production stress. Since the Tr.CBH elicited a stronger UPR response in our previous experiment, we introduced the G-SM1 biosensor and low copy Tr.CBH expression vector into the CEN.PK2-1C strain and evaluated its respective secretion capacity and UPR activation, relative to the BY4742 strain (Figure 6).

**Figure 6.**
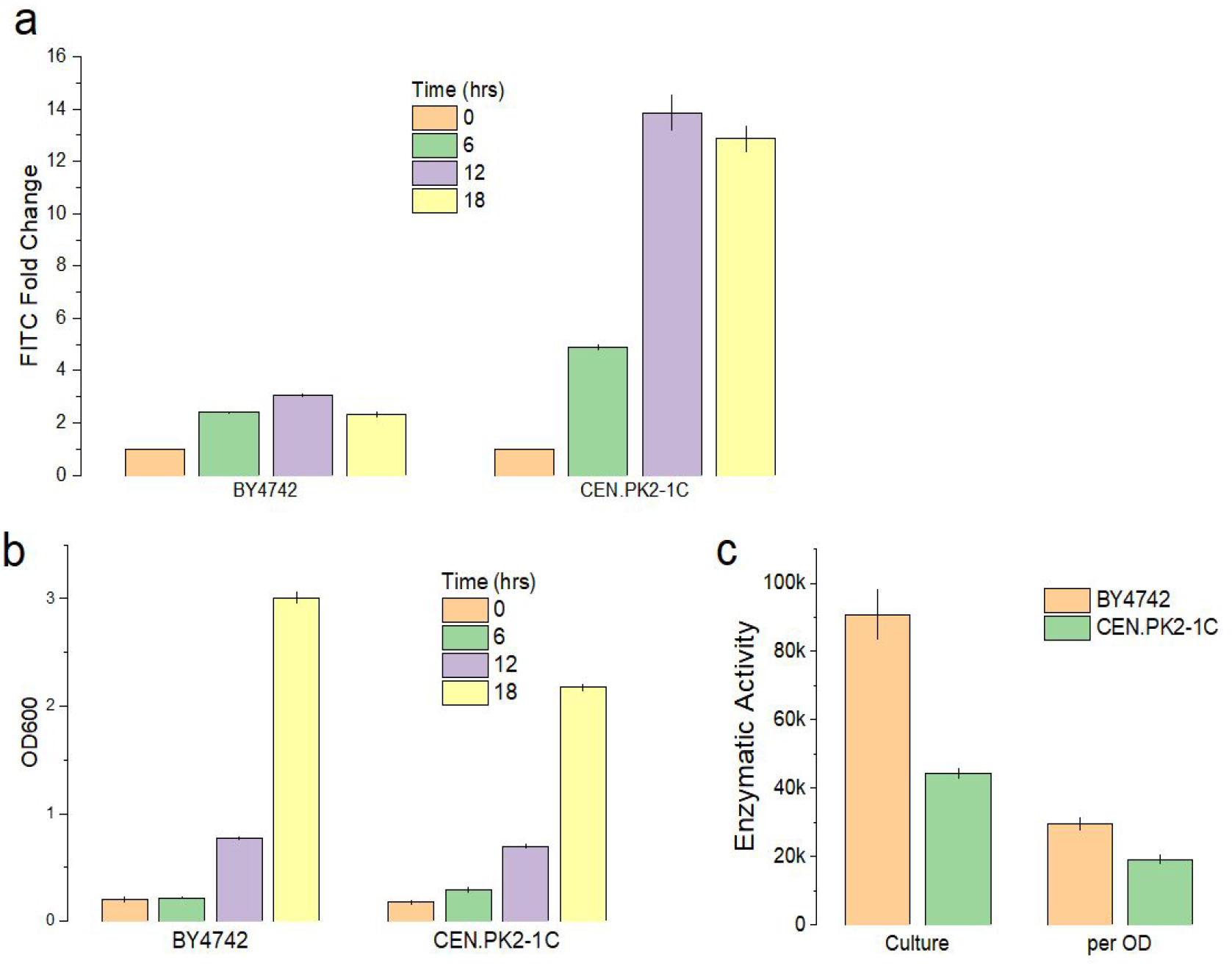
The difference in ER stress tolerance in BY4742 and CEN.PK2-1C strains, and the correlation with Tr.CBH secretion titre, expressed via low copy vector using the two strains as host. (a) The UPR sensor signal was read at 0, 6, 12 and 18 hours after inducing the Tr.CBH expression. (b) The cell density (OD600) measured at each time point. (c) The Tr.CBH activity was compared through MULac assay done as previously. The error bars represent the sample standard deviation from at least three biological replicates.

After 12 hours’ incubation, the G-SM1 sensor response rose to around 3 fold the basal level fluorescence (T_0_) in BY4742 strain consistent with our previous result (Figure 5a). In sharp contrast, the CEN.PK2-1C strains fluorescence signal increased almost 14 folds over the same growth period (Figure 6a). Although both strains have been established as commonly used lab strains, the significantly stronger UPR activation in the CEN.PK2-1C strain demonstrated the implication of the similar, but still heterogeneous genetic background on common stress responses. As expected from our previous data, the strong UPR response of the CEN.PK2-1C strain was also correlated with a lowered extracellular Tr.CBH activity, with the BY4742 culture having over twice the activity by 18 hours (Figure 6c). This echoed the observations that lower UPR leads to higher secretion, and indicated that this UPR/secretion relationship could potentially be exploited using biological sensors to facilitate high-throughput screening to survey the secretion capacities of different *S. cerevisiae* strains.

#### Summary

The optimal production of recombinant proteins in yeast is dependent on the delicate balance between the protein synthesis load and the ER’s ability to effectively process it. This balance is further influenced by external factors such as the growth media composition, the biophysical properties of the protein of interest and the small, but significant differences in cellular metabolism between genetically distinct strains. Here we have constructed and evaluated novel synthetic minimal UPR biosensors with enhanced resolution, dynamic and working ranges. These sensors were modular in design and utilised updated genetic information on the regulatory elements of UPR genes. Considering the tremendous impact of the ER’s folding capability on the cells ability to produce and secrete proteins, having an early indication of stress and its concomitant effect on protein yields has substantial fundamental and industrial value. Here we were able to demonstrate the direct application of these sensors as effective early reporters of UPR and its direct application to predict high and low protein secretion. Our sensor was able to detect UPR differences caused by different proteins, abnormalities in protein folding and expression levels. Although the reasons for many of the highly protein-specific differences in protein production is still unknown, biosensors like the ones described here represent significant advancement in the detection of a major bottleneck in protein processing and opens the door for high-throughput screening approaches.

## Materials and Methods

### Strains and Media

*S. cerevisiae* strain CEN.PK2-1C was used for UPR sensor evaluation using tunicamycin as ER stress inducer. To make the CEN.PK2-1C Δhac strain, PCR fragment was obtained from the genomic DNA of BY4741 strain with *hac1* deletion, and then integrated to the chromosome of CEN.PK2-1C to knockout the *HAC1* gene.

In the experiments involving overexpressed protein as UPR inducer, BY4742 strain was used (*MATalpha his3Δ1 leu2Δ0 met15Δ0 ura3Δ0*). CEN.PK2-1C and BY4742 strains carrying UPR sensors either on plasmids or as single copy chromosomal integrations were constructed.

Mid to late log phase cells were inoculated into 2xSC^−ura^(Glucose) (1.36% w/v yeast nitrogen base, 0.384% w/v yeast dropout supplement without uracil, 2% w/v glucose, 2% w/v succinic acid and 1.2% w/v sodium hydroxide, adjust pH to 6.0) media to obtain starting optical densities of 0.15 at OD_600_ for tests using tunicamycin and the *hac1* deletion strains. And relevant concentration of tunicamycin was added before the experiment. In CPY/CPY* and CBH tests, 2xSC^−his^(Galactose) media (same as 2xSC^−ura^(Glucose) except 0.38% w/v yeast dropout supplement without histidine and 2% galactose instead of glucose were added) was used.

Cells were grown at 30 degrees and 200 RPM. In the native UPR promoter test, the cells were grown in shake flask at a column of 30mL. In the CBH experiment, 5mL of culture were incubated in 50mL Falcon tube sealed with gas-permeable membrane; and in the rest of the study, 1mL culture in 24 well plates were used.

All the yeast transformation was done using the LiOAc/ssDNA method described in Gietz 2007.

### Plasmids and Cloning

The 9 native UPR promoters were PCR amplified from the genomic DNA of CEN.PK2-1C using the primers listed in Table 2S. The PCR products were linked to eGFP and CYC1 terminator, and the resulting sensor cassettes were then cloned into the multi-cloning site of pRS416 vector.

The semi-synthetic UPR promoter and the 4 synthetic minimal UPR promoters were synthesized at GeneScript. The *DER1* and *ERO1* native promoter were obtained as mentioned above. These promoters were built into sensor cassettes as above and cloned into the vector pFGHF_SM1, flanked flo8 homologous sequence and *URA3* marker. PCR products for chromosomally integrated into the CEN.PK2-1C and BY4742 strains. All the UPR sensor constructs were listed in Figure 7. The sequence of yeast synthetic minimal promoter is listed in Table 2S.

**Figure 7.**
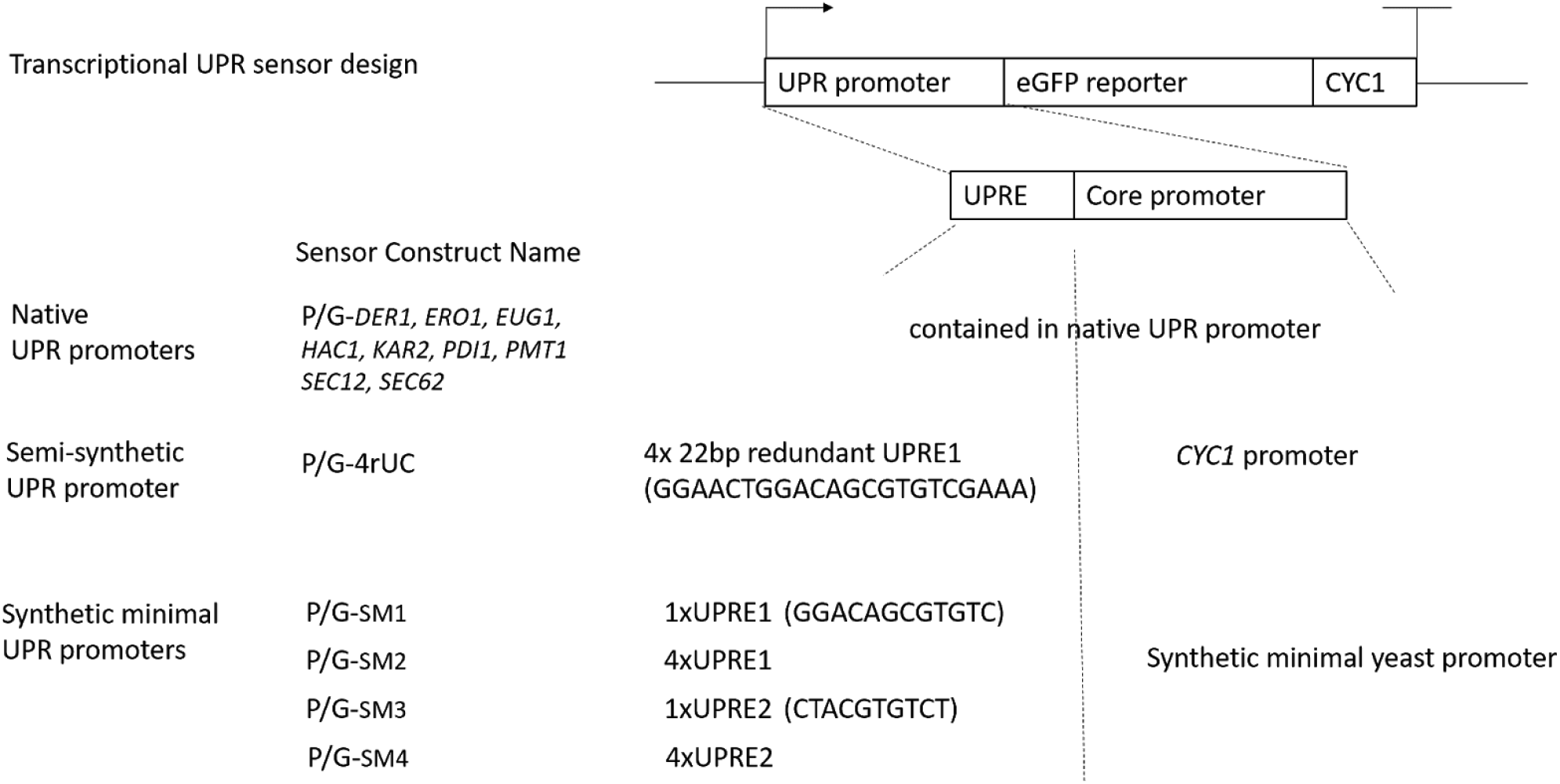
The diagram of UPR sensor design used in this study showing the combination of different UPRE and core promoters. The abbreviation name for each sensor cassette is listed: P- plasmid form UPR sensor; G- chromosome integrated UPR sensor.

The CPY and CPY* cds were amplified from the genomic DNA of CEN.PK2-1C. The Te.CBH and Te.CBH cds were extracted from plasmid pMU(hph)_Te.CBH1 (Kroukamp 2017) via PacI/AscI restriction enzyme digest. All these fragments were built into expression cassettes with *GAL1* promoter and *CYC1* terminator, and subsequently cloned into pRS413 and pRS423 backbone to make the relevant expression vectors. The primers used in this study are listed in Table S2.

### Flow Cytometry

At relevant time point, yeast culture was sampled and loaded on 96-well plates. Dilute to OD600 <=0.5 if necessary. Beckman Cytoflex S was used to fast assay the mean GFP fluorescence of the cells through the FITC channel (ex/em 488/525nm) to record 10000 events. Gating was applied to select the main population (>80% events) if the sample showed discrete FITC distribution (Figure S2).

### Mulac Assay

MULac assays were performed as described by Kroukamp 2017. For each MULac reaction, 25μL substrate in acetate buffer was mixed with 25μL supernatant from cell culture. The reaction mix was incubated in Eppendorf tubes at 42 degrees using a water bath for 30 minutes for Te.CBH, and 4 hours for Tr.CBH. Reaction was stopped by adding 50μL 1M sodium carbonate and the final mix was transferred to black 96-well plate. Fluorescence was measured using a BioTek plate reader with ex/em wavelength at 305/420nm. The 2xSC^−his^(Galactose) media was used in place of culture supernatant following the above preparation as blank.

## References

[1] Mori, K. (2009) Signalling pathways in the unfolded protein response: development from yeast to mammals, J Biochem 146, 743–750.

[2] Walter, P., and Ron, D. (2011) The unfolded protein response: from stress pathway to homeostatic regulation, Science 334, 1081–1086.

[3] Lajoie, P., Moir, R. D., Willis, I. M., and Snapp, E. L. (2012) Kar2p availability defines distinct forms of endoplasmic reticulum stress in living cells, Mol Biol Cell 23, 955–964.

[4] Promlek, T., Ishiwata-Kimata, Y., Shido, M., Sakuramoto, M., Kohno, K., and Kimata, Y. (2011) Membrane aberrancy and unfolded proteins activate the endoplasmic reticulum stress sensor Ire1 in different ways, Mol Biol Cell 22, 3520–3532.

[5] Volmer, R., van der Ploeg, K., and Ron, D. (2013) Membrane lipid saturation activates endoplasmic reticulum unfolded protein response transducers through their transmembrane domains, Proceedings of the National Academy of Sciences 110, 4628–4633.

[6] Volmer, R., and Ron, D. (2015) Lipid-dependent regulation of the unfolded protein response, Curr Opin Cell Biol 33, 67–73.

[7] Kimata, Y., Ishiwata-Kimata, Y., Yamada, S., and Kohno, K. (2006) Yeast unfolded protein response pathway regulates expression of genes for anti-oxidative stress and for cell surface proteins, Genes Cells 11, 59–69.

[8] Bernales, S., McDonald, K. L., and Walter, P. (2006) Autophagy counterbalances endoplasmic reticulum expansion during the unfolded protein response, PLoS Biol 4, e423.

[9] Kevin J Travers, C. K. P., Lisa Wodicka, David J Lockhart, Jonathan S Weissman, Peter Walter. (2000) Functional and Genomic Analyses Reveal an Essential Coordination between the Unfolded Protein Response and ER-Associated Degradation, Cell 101, 249–258.

[10] Ng, D. T., Spear, E. D., and Walter, P. (2000) The unfolded protein response regulates multiple aspects of secretory and membrane protein biogenesis and endoplasmic reticulum quality control, J Cell Biol 150, 77–88.

[11] Hou, J., Tyo, K., Liu, Z., Petranovic, D., and Nielsen, J. (2012) Metabolic engineering of recombinant protein secretion by Saccharomyces cerevisiae. FEMS Yeast Research 12, 491–510.

[12] Ma, Y., and Hendershot, L. M. (2004) The role of the unfolded protein response in tumour development: friend or foe?, Nat Rev Cancer 4, 966–977.

[13] Lin, J., Li, H., Yasumura, D., Cohen, H., Zhang, C., Panning, B., Shokat, K., LaVail, M., and Walter, P. (2007) IRE1 Signaling Affects Cell Fate During the Unfolded Protein Response. Science 318, 944–949.

[14] Cox, J., and Walter, P. (1996) A Novel Mechanism for Regulating Activity of a Transcription Factor That Controls the Unfolded Protein Response. Cell 87, 391–404.

[15] Pincus, D., Chevalier, M. W., Aragon, T., van Anken, E., Vidal, S. E., El-Samad, H., and Walter, P. (2010) BiP binding to the ER-stress sensor Ire1 tunes the homeostatic behavior of the unfolded protein response, PLoS Biol 8, e1000415.

[16] Gardner, B. M., & Walter, P. (2011). Unfolded proteins are Ire1-activating ligands that directly induce the unfolded protein response. Science (New York, N.Y.), 333(6051), 1891–1894.

[17] Korennykh, A., Egea, P., Korostelev, A., Finer-Moore, J., Zhang, C., Shokat, K., Stroud, R., and Walter, P. (2009) The unfolded protein response signals through high-order assembly of Ire1. Nature 457, 687–693.

[18] van Anken, E., Pincus, D., Coyle, S., Aragon, T., Osman, C., Lari, F., Gomez Puerta, S., Korennykh, A. V., and Walter, P. (2014) Specificity in endoplasmic reticulum-stress signaling in yeast entails a step-wise engagement of HAC1 mRNA to clusters of the stress sensor Ire1, Elife 3, e05031.

[19] Sidrauski, C., & Walter, P. (1997). The transmembrane kinase Ire1p is a site-specific endonuclease that initiates mRNA splicing in the unfolded protein response. Cell, 90(6), 1031–1039. https://doi.org/10.1016/s0092-8674(00)80369-4

[20] Mori, K., Sant, A., Kohno, K., Normington, K., Gething, M., and Sambrook, J. (1992) A 22 bp cis-acting element is necessary and sufficient for the induction of the yeast KAR2 (BiP) gene by unfolded proteins. The EMBO Journal 11, 2583–2593.

[21] Mori, K., Ogawa, N., Kawahara, T., Yanagi, H., & Yura, T. (1998). Palindrome with spacer of one nucleotide is characteristic of the cis-acting unfolded protein response element in Saccharomyces cerevisiae. The Journal of biological chemistry, 273(16), 9912–9920. https://doi.org/10.1074/jbc.273.16.9912

[22] Fordyce, P. M., Pincus, D., Kimmig, P., Nelson, C. S., El-Samad, H., Walter, P., and DeRisi, J. L. (2012) Basic leucine zipper transcription factor Hac1 binds DNA in two distinct modes as revealed by microfluidic analyses, Proceedings of the National Academy of Sciences 109, E3084–E3093.

[23] Gasser, B., Saloheimo, M., Rinas, U., Dragosits, M., Rodriguez-Carmona, E., Baumann, K., Giuliani, M., Parrilli, E., Branduardi, P., Lang, C., Porro, D., Ferrer, P., Tutino, M. L., Mattanovich, D., and Villaverde, A. (2008) Protein folding and conformational stress in microbial cells producing recombinant proteins: a host comparative overview, Microb Cell Fact 7, 11.

[24] Xu, L., Shen, Y., Hou, J., Peng, B., Tang, H., and Bao, X. (2014) Secretory pathway engineering enhances secretion of cellobiohydrolase I from Trichoderma reesei in Saccharomyces cerevisiae. Journal of Bioscience and Bioengineering 117, 45–52.

[25] Merksamer, P. I., Trusina, A., and Papa, F. R. (2008) Real-time redox measurements during endoplasmic reticulum stress reveal interlinked protein folding functions, Cell 135, 933–947.

[26] Le, Q. G., Ishiwata-Kimata, Y., Kohno, K., and Kimata, Y. (2016) Cadmium impairs protein folding in the endoplasmic reticulum and induces the unfolded protein response, FEMS Yeast Res 16.

[27] Patil, C., Li, H., and Walter, P. (2004) Gcn4p and Novel Upstream Activating Sequences Regulate Targets of the Unfolded Protein Response. PLoS Biology 2, e246.

[28] Xu, P., Raden, D., Doyle, F., and Robinson, A. (2005) Analysis of unfolded protein response during single-chain antibody expression in Saccharomyces cerevisiae reveals different roles for BiP and PDI in folding. Metabolic Engineering 7, 269–279.

[29] Aragon, T., van Anken, E., Pincus, D., Serafimova, I. M., Korennykh, A. V., Rubio, C. A., and Walter, P. (2009) Messenger RNA targeting to endoplasmic reticulum stress signalling sites, Nature 457, 736–740.

[30] Cedras, G., Kroukamp, H., Van Zyl, W. H., and Den Haan, R. (2020) The in vivo detection and measurement of the unfolded protein response in recombinant cellulase producing Saccharomyces cerevisiae strains, Biotechnol Appl Biochem 67, 82–94.

[31] Redden, H., and Alper, H. S. (2015) The development and characterization of synthetic minimal yeast promoters, Nat Commun 6, 7810.

[32] de Ruijter, J. C., Koskela, E. V., Nandania, J., Frey, A. D., and Velagapudi, V. (2018) Understanding the metabolic burden of recombinant antibody production in Saccharomyces cerevisiae using a quantitative metabolomics approach, Yeast 35, 331–341.

[33] Finger, A., Knop, M., and Wolf, D. H. (1993) Analysis of two mutated vacuolar proteins reveals a degradation pathway in the endoplasmic reticulum or a related compartment of yeast, Eur J Biochem 218, 565–574.

[34] Kuhad, R., Gupta, R., and Singh, A. (2011) Microbial Cellulases and Their Industrial Applications. Enzyme Research 2011, 1–10.

[35] Ilmén, M., den Haan, R., Brevnova, E., McBride, J., Wiswall, E., Froehlich, A., Koivula, A., Voutilainen, S., Siika-aho, M., la Grange, D., Thorngren, N., Ahlgren, S., Mellon, M., Deleault, K., Rajgarhia, V., van Zyl, W., and Penttilä, M. (2011) High level secretion of cellobiohydrolases by Saccharomyces cerevisiae. Biotechnology for Biofuels 4, 30.

[36] Gorgens, J. F., van Zyl, W. H., Knoetze, J. H., and Hahn-Hagerdal, B. (2001) The metabolic burden of the PGK1 and ADH2 promoter systems for heterologous xylanase production by *Saccharomyces cerevisiae* in defined medium. Biotechnol Bioeng 73, 238–245.

[37] Kroukamp, H., den Haan, R., la Grange, D. C., Sibanda, N., Foulquié-Moreno, M. R., Thevelein, J. M., van Zyl, W. H. (2017) Strain breeding enhanced heterologous cellobiohydrolase secretion by *Saccharomyces cerevisiae* in a protein specific manner. Biotechnol J 12:1700346

[38] Valkonen, M., Penttilä, M., and Saloheimo, M. (2003) Effects of Inactivation and Constitutive Expression of the Unfolded-Protein Response Pathway on Protein Production in the Yeast *Saccharomyces cerevisiae*. Appl Environ Microbiol 69, 2065–2072.

[39] Kroukamp, H., den Haan, R., van Zyl, J-H., van Zyl, W. H. (2018) Rational strain engineering interventions to enhance cellulase secretion by *Saccharomyces cerevisiae*. Biofuel Bioprod Biorefin 12:108–124

[40] Kroukamp, H., den Haan, R., van Wyk, N., van Zyl, W. H. (2013) Overexpression of native *PSE1* and *SOD1* in *Saccharomyces cerevisiae* improved heterologous cellulase secretion

