## Supplementary data for "Yeast synthetic minimal biosensors for evaluating protein production"

### Summary

Table 1S. Strains used in this study

| Strain | Description |
|---|---|
| CEN.PK2-1C | Wildtype |
| BY4741 del hac | Δhac1::KanMX |
| CEN-Δhac1 | Δhac1::KanMX |
| BY4742 | Wildtype |
| BY42-SM1 | Δflo8::URA3 with integrated SM1 UPR sensor |

Table 2S. The nucleotide sequence of the 4 synthetic minimal UPR promoters.

| Promoters | Sequence |
|---|---|
| SM1 | GGACAGCGTGTCCTTAAGATCTTGTAATATTCTAATCAAGCTTATAAAAGAGCACTGTTGGGCGTGAGTGGAGGCGCCGGAATAAACCATGCAATGAA |
| SM2 | GGACAGCGTGTCGGACAGCGTGTCGGACAGCGTGTCGGACAGCGTGTCCTTAAGATCTTGTAATATTCTAATCAAGCTTATAAAAGAGCACTGTTGGGCGTGAGTGGAGGCGCCGGAATAAACCATGCAATGAA |
| SM3 | CTACGTGTCACTTAAGATCTTGTAATATTCTAATCAAGCTTATAAAAGAGCACTGTTGGGCGTGAGTGGAGGCGCCGGAATAAACCATGCAATGAA |
| SM4 | CTACGTGTCACTACGTGTCACTACGTGTCACTACGTGTCACTTAAGATCTTGTAATATTCTAATCAAGCTTATAAAAGAGCACTGTTGGGCGTGAGTGGAGGCGCCGGAATAAACCATGCAATGAA |

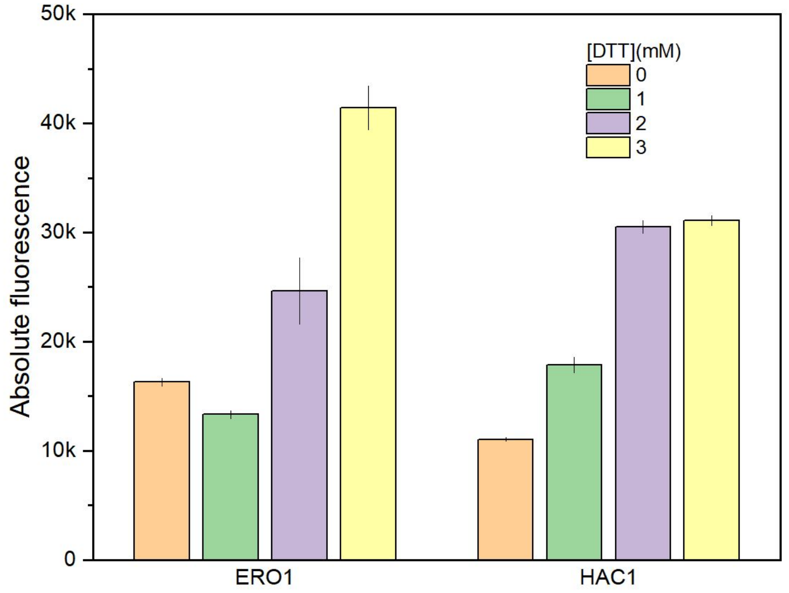


Figure 1S. The response of UPR sensors constructed with P-ERO1 and P-HAC1 promoters, showing the insensitivity of P-HAC1 between 2mM and 3mM DTT. Cells were incubated in DTT 4 hours. Each point represented 3 biological replicates.

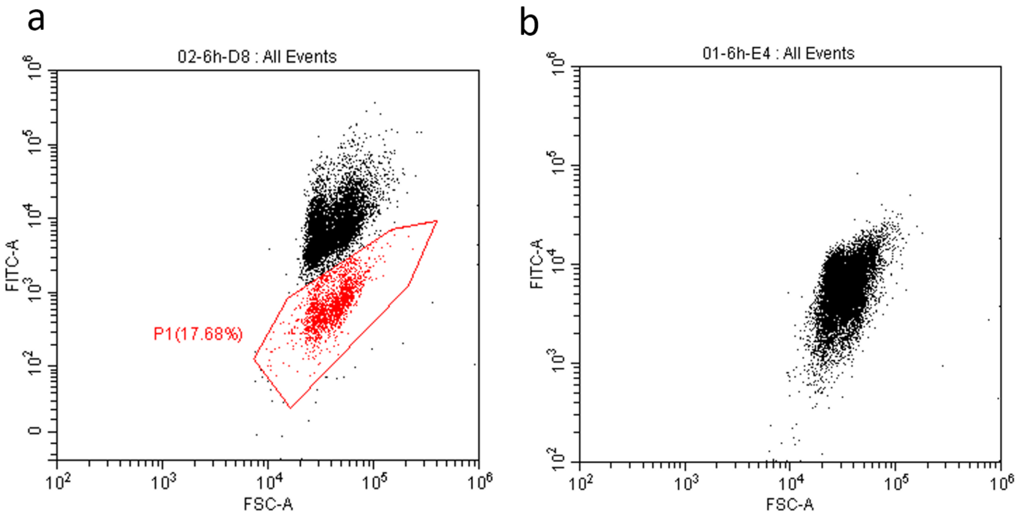


Figure 2S. The fluorescence-plot of the SM1 sensor as plasmid form P-SM1 (a) and chromosomally integrated form G-SM1 (b). The noise subpopulation was indicated in (a), which diminished in (b), showing the improvement in the FITC population homogeneity after chromosome integration. Cells were treated with 0.5 μg/mL Tm for 6 hours.
